# Challenging a role for ceramide channels and microdomains in apoptosis induction using a bottom-up approach

**DOI:** 10.1101/2025.11.14.688508

**Authors:** Milena Wessing, Sophie Klaßen, Britta Fiedler, Katia Cosentino, Joost C. M. Holthuis

## Abstract

Ceramides are essential but potentially toxic intermediates of sphingolipid metabolism that can act directly on mitochondria to trigger apoptotic cell death, but the underlying mechanism is unclear. While one model postulates that ceramides form stable channels in the outer mitochondrial membrane that induce cell death through direct release of cytochrome c, an alternative view is that ceramides self-assemble into microdomains that facilitate membrane insertion and oligomerization of the pro-apoptotic Bcl-2 protein Bax into cytochrome *c*-conducting pores. To challenge these models, we here analyzed the influence of ceramides in combination with recombinant Bax on the leakiness of model membranes. We show that ceramides on their own are unable to support membrane passage of even the smallest fluorescence markers. Moreover, we find that ceramides cannot substitute for cardiolipin in facilitating membrane recruitment of Bax and its subsequent assembly into functional pores. Our data argue against a direct role of ceramides in apoptotic pore formation and indicate that the mechanism by which ceramides initiate permeabilization of the outer mitochondrial membrane is independent of ceramide channels or ceramide acting autonomously as translocation platform for Bax.

## INTRODUCTION

Apoptosis is the best understood form of regulated cell death and plays an essential role in organismal development, tissue homeostasis and a correctly functioning immune system. Perturbations in apoptosis contribute to human disease, notably cancer and autoimmunity [1]. Apoptosis induction by stress stimuli (e.g. DNA damage, cytokines or cytostatics) converges on mitochondria, whereby the opening of apoptotic pores in the outer mitochondrial membrane [2] and cytosolic release of cytochrome *c* defines a point of no return, leading to caspase activation and the ordered self-destruction of the cell [3]. OMM permeabilization for cytochrome *c* is tightly controlled by the B-cell lymphoma 2 (Bcl-2) protein family, which includes pro- and anti-apoptotic members that collectively determine the balance between cell death and survival [4]. The principal pro-apoptotic Bcl-2 proteins in mammalian cells are Bax and Bak. In healthy cells, while Bak mainly resides at mitochondria, Bax constitutively cycles between mitochondria and the cytosol due to the action of anti-apoptotic Bcl-2 family members [5]. In response to DNA damage or other forms of cellular stress, Bax accumulates on the mitochondrial surface, where it undergoes extensive conformational rearrangements and oligomerization to form cytochrome *c*-conducting pores [6–8].

While research on the mechanisms underlying OMM permeabilization in mitochondrial apoptosis primarily focuses on the role of Bcl-2 proteins, there are also reports that emphasize a crucial role for lipids. Notably ceramides, the direct precursors of sphingomyelin and glycosphingolipids, have frequently been implicated as potent mediators of stress-induced mitochondrial apoptosis [9]. A rise in mitochondrial ceramide levels has frequently been observed to precede OMM permeabilization in response to a variety of distinct stresses, including exposure to ionizing radiation, tumor necrosis factor TNFa, and chemotherapeutics [10–13]. Suppression of ceramide accumulation renders cells resistant to these apoptotic stimuli, supporting a direct role of ceramides in stress-induced apoptotic cell death. Numerous studies revealed that ceramides can act directly on mitochondria to initiate permeabilization of the OMM for cytochrome *c* [9]. For instance, mitochondrial targeting of a bacterial sphingomyelinase to generate ceramides in mitochondria induces cytochrome *c* release and apoptosis [14]. Moreover, diverting CERT-mediated ceramide transport to mitochondria triggers Bax-dependent apoptosis [15, 16]. Apoptosis induction is abolished by mitochondrial targeting of a bacterial ceramidase, indicating that apoptogenic activity relies on intact ceramides rather than metabolic intermediates of ceramide turnover. The notion that ceramides can initiate apoptotic cell death by acting directly on mitochondria was recently reinforced by the observation that mitochondria-specific photorelease of ceramides triggers cleavage of Caspase 9, signifying activation of the intrinsic apoptotic pathway [17].

How a rise in mitochondrial ceramides initiates OMM permeabilization to commit cells to death is subject of debate. One model postulates that ceramides can self-assemble into channels in the OMM that are large enough to facilitate passage of small proteins including cytochrome *c* [18]. This concept was first described in a study reporting that ceramides induce vertical increments in current across planar bilayers under voltage clamp conditions in the absence of proteins [19, 20]. Molecular dynamics simulations revealed that the structural units of ceramide channels are columns of four to six ceramides that are H-bonded via amide groups and arranged as staves in either a parallel or antiparallel manner [21]. Based on the conductance of single channels [19, 20], the molecular weight of proteins released from ceramide-treated mitochondria [22, 23] and visualization of pores in ceramide-containing liposomes by negative stain electron microscopy [24], a typical ceramide channel was estimated to have a pore diameter of approximately 10 nm. The pore-forming activity of ceramides did not require any auxiliary proteins but was directly inhibited by anti-apoptotic Bcl-2 proteins [25, 26]. However, proof that ceramide channels mediate OMM permeabilization *in vivo* is lacking. Moreover, the results of a study addressing ceramide-mediated leakage in sphingomyelin-rich liposomes argued against ceramide channels as the mechanism for ceramide-induced membrane permeabilization [27].

An alternative model postulates that mitochondrial ceramides self-assemble into microdomains that facilitate functionalization of Bax to form cytochrome *c*-conducting pores. This notion originated from studies demonstrating that mitochondrial ceramide generation is obligate for radiation-induced apoptosis in the *C. elegans* germline [13]. Furthermore, in isolated mitochondria exogenously added ceramides and recombinant Bax acted coordinately to release cytochrome *c* [28, 29]. A subsequent study revealed that ceramides generated in the OMM of mammalian cells upon irradiation formed microdomains that could be visualized and isolated biophysically, into which Bax integrated [30]. Pharmacological inhibition of ceramide generation prevented radiation-induced Bax insertion, oligomerization and OMM permeabilization. Other studies with pore-forming toxins and recombinant Bax indicated that pore assembly involves the formation of a toroidal structure, where pore walls interact directly with lipids such that non-bilayer structures are generated, enabling pore opening [31]. Ceramides belong to one of only few lipid classes that have the propensity to spontaneously “destabilize” membranes by hexagonal phase transition [32, 33].

Consequently, ceramide microdomains formed in the OMM of irradiated cells may function as platforms into which Bax inserts, oligomerizes and converts into an open, functional pore [30].

To shed further light on the mechanism by which ceramides commit cells to death, we here pursued a bottom-up approach in which we analyzed the influence of ceramides in combination with Bax on the leakiness of model membranes for fluorescence size markers. We find that ceramides on their own are unable to support membrane passage of even the smallest fluorescence markers. Additionally, we find that ceramides cannot substitute for cardiolipin in facilitating membrane recruitment of Bax and its subsequent assembly into pores that mediate the release of fluorescence markers in the size range of cytochrome *c*. Our data argue against a direct role of ceramides in apoptotic pore formation and indicate that the mechanism by which ceramides initiate permeabilization of the OMM is independent of ceramide channels or ceramide acting as translocation platform for Bax.

## RESULTS

### Production and functional characterization of recombinant Bax

To obtain functionally competent, monomeric and fluorescently labeled recombinant Bax, we used an IPTG-inducible bacterial expression construct encoding a human Bax single cysteine mutant (S4C, C62S, C126S) tagged with a *C*-terminal intein–chitin binding domain (CBD; **Fig. 1a**). Presence of the intein-CBD tag enabled affinity purification of the fusion protein on a chitin column. A DTT-induced self-cleavage of intein released Bax from the chitin beads, resulting in an eluate containing full-length recombinant Bax with no exogenous residues (**Fig. 1b**). Subsequent anion exchange chromatography enabled removal of DTT and residual contaminating bacterial proteins (**Fig. 1c**). Next, purified recombinant Bax was subjected to cysteine labeling with DY-647P1. Size exclusion chromatography resulted in an efficient separation of monomeric, fluorescently labeled Bax from Bax dimers and free DY-647P1 (**Fig. 1d**). The degree of Bax labeling was ∼74 %. Final concentrations were 10 µM for the unlabeled and 6.25 µM for the labeled Bax.

**Figure 1.**
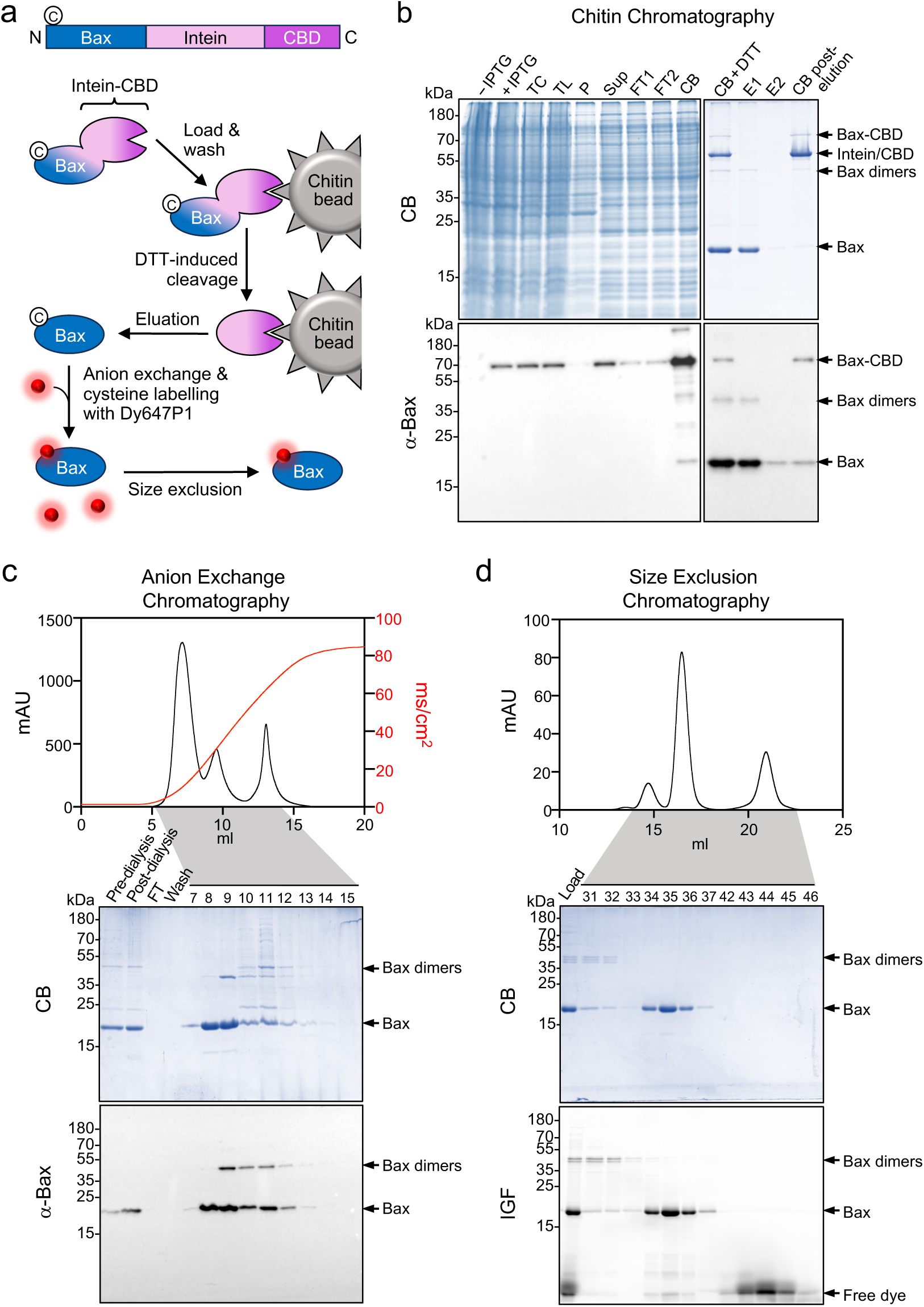
Purification and cysteine labelling of recombinant Bax. **(a)** Schematic outline of the production, purification and cysteine-labeling of recombinant human Bax using an IPTG-inducible bacterial expression construct encoding the Bax single cysteine mutant (S4C, C62S, C126S) tagged with a *C*-terminal intein-chitin binding domain (CBD). **(b)** Chitin affinity chromatography of the Bax-intein-CBD fusion protein upon its IPTG-induced expression in *E. coli*. After binding of the fusion protein to chitin resin, native Bax was released by DTT-induced self-cleavage of intein. Expression and proteolytic release of Bax was monitored by SDS-PAGE, Coomassie Blue (CB) staining and immunoblot (IB) analysis using an αBax antibody. TC, total cells; TL, total lysate; P, pellet; Sup, supernatant; FT, flow through; CB, chitin beads. **(c)** Bax collected by chitin affinity chromatography was dialyzed and loaded onto an anion exchange column concentrated and Anion exchange chromatography of recombinant. fractions recombinant Bax. To reach a higher purity of the protein Bax was dialyzed and loaded onto an anion exchange column that was developed with a linear salt gradient. Peak fractions were analyzed by SDS-PAGE, CB staining and IB analysis as in (b). **(d)** Bax collected by anion exchange chromatography was cysteine-labeled with DY-647P1 and loaded onto a size exclusion column. Peak fractions were analyzed by SDS-PAGE, CB staining and in-gel-fluorescence (IGF) analysis.

To validate that recombinant DY-647P1-labeled Bax was functional and retained its pore-forming activity, we conducted a calcein-release assay (**Fig. 2a**). Detection of calcein leakage from large unilamellar vesicles (LUVs) is an effective indicator of membrane permeabilization, as calcein undergoes self-quenching at high concentrations inside the vesicles but displays fluorescence emission upon dilution in the surrounding buffer [34, 35]. Accordingly, LUVs enclosing calcein at a self-quenching concentration were exposed to either unlabeled or DY-647P1-labeled Bax. Caspase 8-cleaved Bid (cBid) was added, which activates Bax to enable its membrane association and self-assembly into a membrane permeabilizing pore [36]. LUVs were prepared from a mixture of eggPC and cardiolipin (CL) at an 80:20 molar ratio, as the presence of CL is critical for an efficient pore forming activity of Bax in model membranes [37, 38]. We initially monitored calcein release kinetics from LUVs in the presence of cBid across a wide range of Bax concentrations (**Fig. 2b**). Calcein leakage was already detectable upon external addition of 10 nM cBid-treated Bax. Calcein release kinetics were enhanced by increasing the concentration of activated Bax. The pore-forming activity of DY-647P1-labeled Bax was only slightly reduced in comparison to that of unlabeled Bax (**Fig. 2b**). In line with the literature [6, 39, 40], omission of either Bax or CL abolished calcein release whereas omission of cBid treatment drastically reduced Bax-mediated leakage of calcein (**Fig. 2c**). The latter finding indicated that a minor fraction of purified recombinant Bax is auto-active and able to form pores without prior exposure to cBid. From this we conclude that DY-647P1-labeled recombinant Bax is functional and capable of forming calcein-conducting pores in CL-containing membrane bilayers.

**Figure 2.**
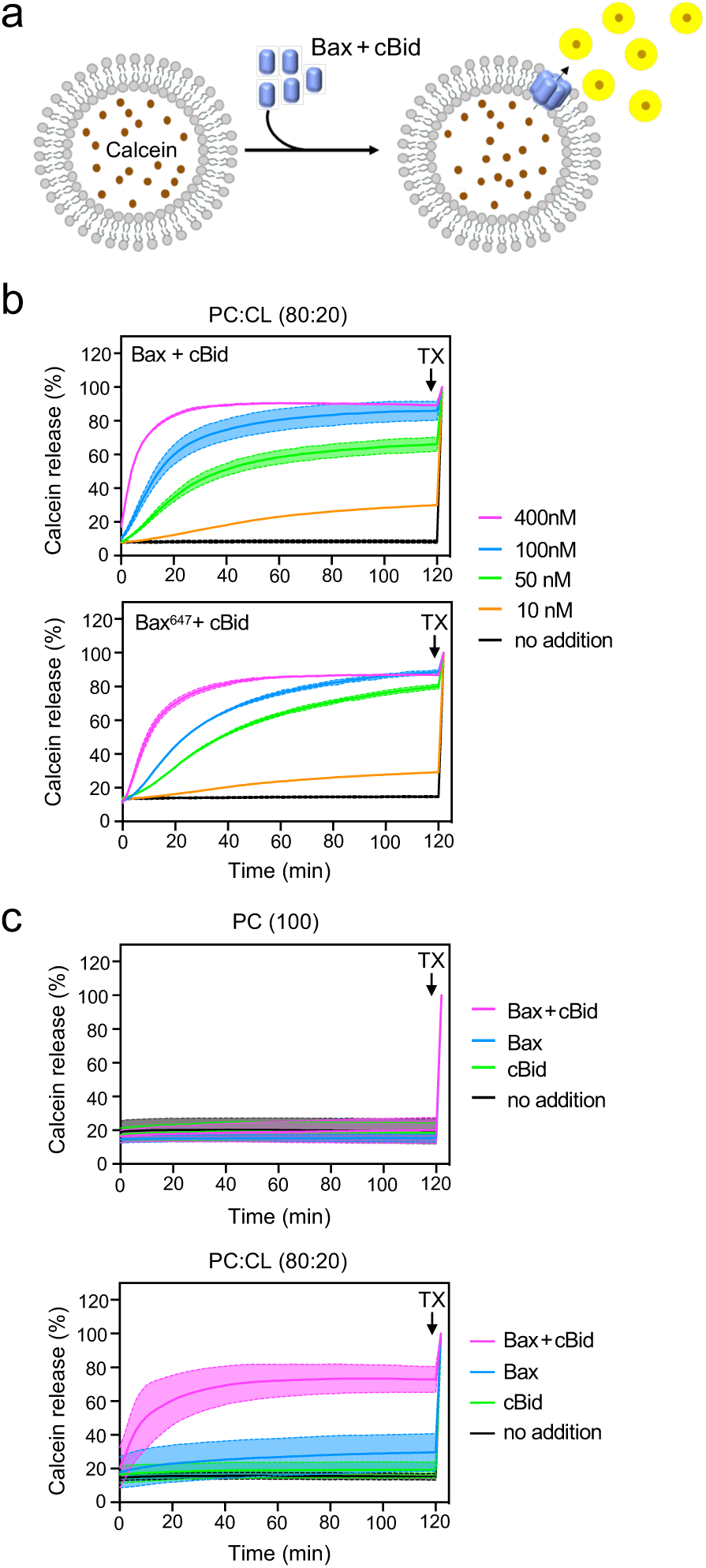
Functional characterization of recombinant Bax. **(a)** Schematic outline of calcein release assay for measuring pore-forming activity of cBid-activated recombinant Bax in large unilamellar vesicles. **(b)** Real-time chromatograms of calcein released from PC:CL (80:20) vesicles co-incubated with the indicated amount of unlabeled Bax or DY-647P1-labeled Bax and 50 nM cBid. Leakiness of vesicles was tested by omitting Bax and cBid (no addition). **(c)** Real-time chromatograms of calcein released from PC (100) and PC:CL (80:20) vesicles incubated with different combinations of 100 nM Bax and 50 nM cBid. Calcein release was measured over 120 min at 2 min intervals using a fluorescence plate reader. For each time point, the mean values of three independent measurements were determined and normalized to the value measured after Triton X100 (TX) addition, which served as a reference for 100% permeabilization.

### Ceramide on its own is unable to create stable pores in model membranes

Both activated Bax and C16:0 ceramide (Cer_16_) have been proposed to form stable pores that are large enough to release 12-kDa cytochrome *c* from mitochondria during apoptosis [7, 18, 19, 43]. This led us to visualize membrane passage of fluorescent size markers across putative pores formed by Bax or Cer_16_ in model membranes using confocal microscopy as described previously [44]. To this end, we generated giant unilamellar vesicles (GUVs) composed of different lipid mixtures and analyzed their leakiness for calcein (0.5 kDa) upon external addition of cBid-activated Bax^647^ (**Fig. 3a**). To enable their visualization in real time, GUVs were stained with lipophilic dyes DiI or DiO. As shown in **Fig. 3b**, external addition of cBid-activated Bax^647^ readily permeabilized CL-containing GUVs for calcein. Leakage of calcein was accompanied by a gradual accumulation of Bax^647^ on the GUV surface and its translocation into the GUV lumen (**Fig. 3b, d**). Omission of CL abolished Bax-mediated calcein leakage and eliminated Bax^647^ binding to the GUV surface as well as its translocation across the GUV bilayer (**Fig. 3b-d**). Moreover, CL-containing GUVs were largely impermeable for calcein in the absence of Bax, even though we observed that GUVs containing 20 mol% CL were generally leakier for calcein than GUVs composed of pure egg PC (**Fig. 3c, Suppl. Fig. 1**). Collectively, these results indicate that in CL-containing GUVs, cBid-activated Bax^647^ self-assembles into pores large enough to allow passage of both calcein (0.5 kDa) and itself (21 kDa) into the GUV lumen.

**Figure 3.**
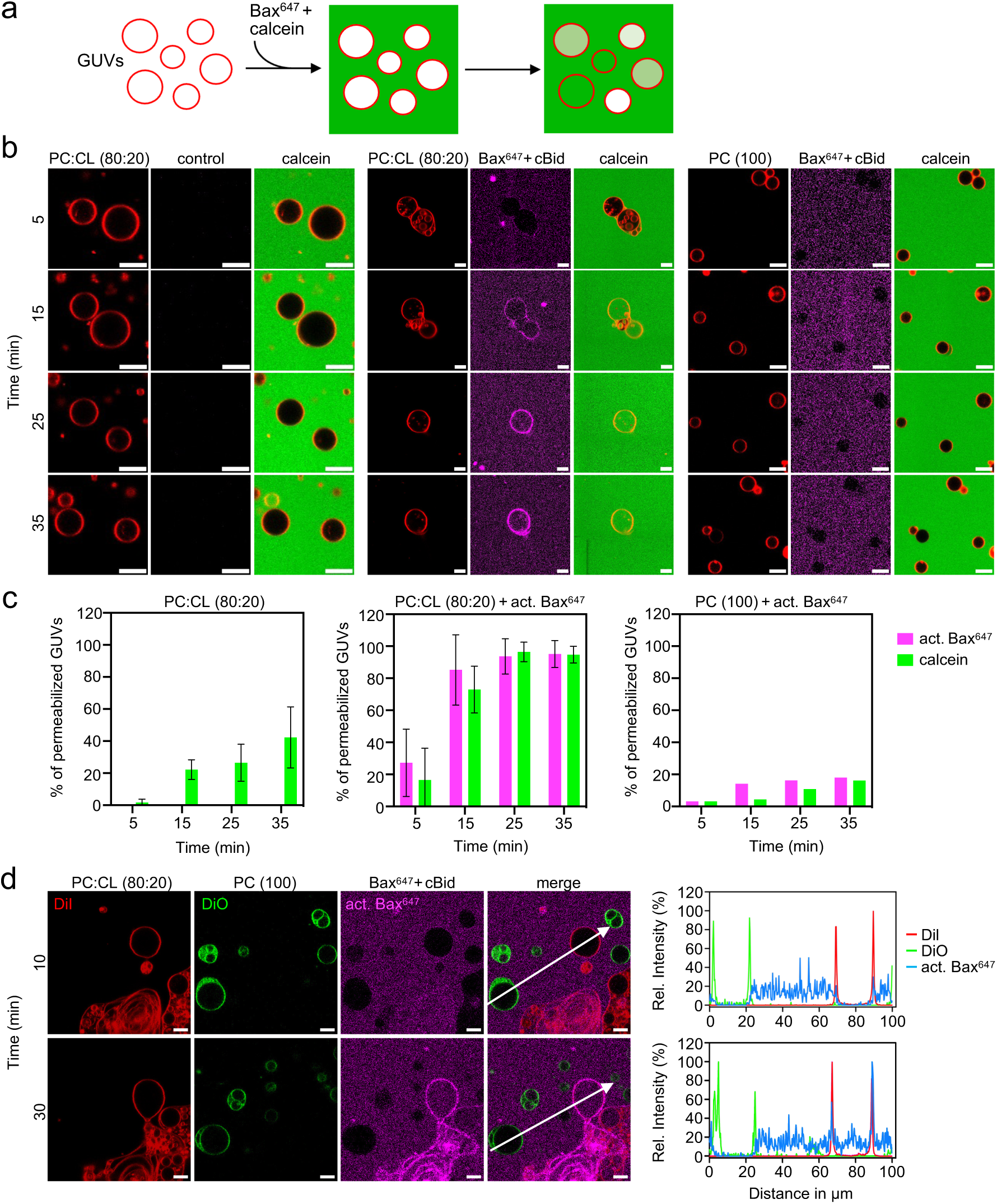
Cardiolipin-mediated Bax recruitment is tightly coupled to membrane permeabilization. **(a)** Schematic outline of calcein permeabilization assay for measuring pore-forming activity of recombinant Bax in giant unilamellar vesicles (GUVs). **(b)** GUVs prepared from PC (100) or PC:CL (80:20) were stained with DiI (*red*) and incubated with calcein (*green*) in the absence or presence of 400 nM Bax^647^ (*magenta*) and 50 nM cBid. GUVs were imaged at the indicated time points using a LSM Airyscan microscope. Scale bar 10 µm. **(c)** Quantification of the permeability of GUVs for Bax^647^(*magenta*) and calcein (*green*) at the indicated incubation times. GUVs with a degree of filling of 40 % or more were considered permeable. For GUVs prepared from PC:CL (80:20), a total number of 70-100 GUVs were analyzed in three independent experiments. For GUVs prepared from PC (100), a total of 50 GUVs were analyzed in one experiment. Data are means ± SD. **(d)** GUVs prepared from PC (100; DiO; *green*) or PC:CL (80:20; DiI; *red*) were incubated in the presence of 400 nM Bax^647^ (*magenta*) and 50 nM cBid. GUVs were imaged as in (b). Line scans along the path of the arrows show the degree of leakage of Bax^647^ inside PC and PC:CL-containing GUVs. Scale bar 10 µm.

To reduce CL-mediated leakiness of GUVs, subsequent experiments were carried out using GUVs with a reduced (10 mol%) CL content. Moreover, Alexa-fluor647 10kDa dextran (Dex10) and fluorescein 70 kDa dextran (Dex70) were used as fluorescent size markers to enable a more accurate assessment of pore sizes. GUVs composed of egg PC or egg PC with 10 mol% CL were impermeable for both Dex10 and Dex70 (**Fig. 4a, b**). As expected, addition of cBid-activated Bax to the CL-containing GUVs caused their gradual permeabilization for both Dex10 and Dex70, with membrane passage of Dex10 occurring with a higher speed and efficiency compared to that of Dex70 (**Fig. 4c**). Substitution of CL for C16:0 ceramide (Cer_16_) or brain ceramide, which predominantly comprises C18:0 ceramide, in each case did not affect the permeability of GUVs for either Dex10 or Dex70 (**Fig. 4d, e**). These results indicate that ceramides on their own are unable to form stable pores in synthetic bilayers that allow passage of polar molecules in the size range of cytochrome *c*.

**Figure 4.**
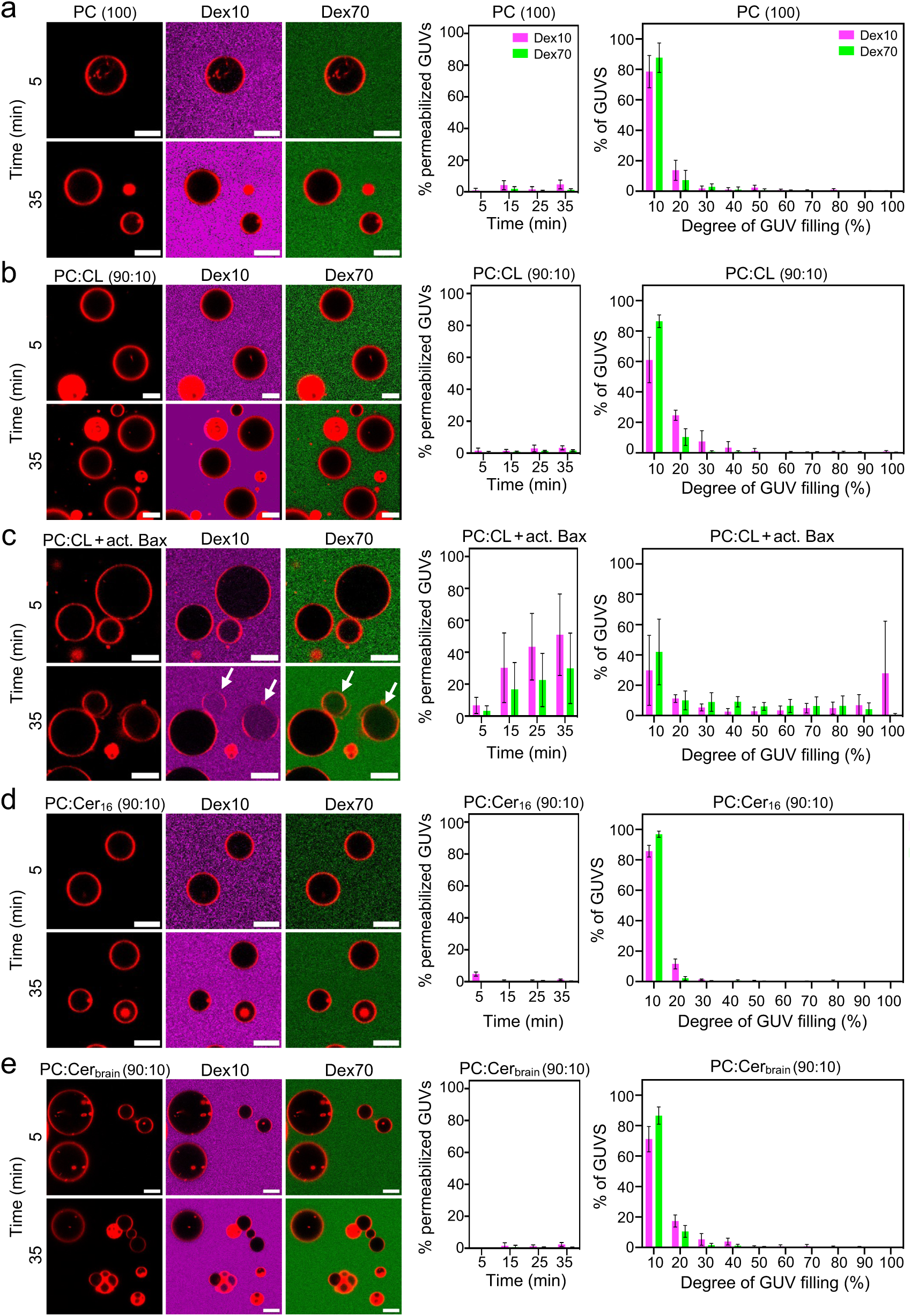
Ceramide-containing model membranes are impermeable for low- and high-molecular weight dextrans. **(a)** GUVs prepared with egg PC (100) and stained with DiI (*red*) were incubated with Alexa-fluor647 10kDa dextran (Dex10; *magenta*) and fluorescein 70 kDa dextran (Dex70; *green*) and then imaged using a LSM Airy scan microscope. At the indicated incubation time, the permeability of the GUVs for Dex10 and Dex70 was quantified. GUVs with a filling degree of 40% or more were considered permeable. In addition, the filling degree of the GUVs was determined at 35 min of incubation. **(b)** GUVs prepared with PC:CL (90:10) were processed as in (a). **(c)** GUVs prepared with PC:CL (90:10) were incubated in the presence of 400 nM Bax and 50 nM cBid and processed as in (a). **(d)** GUVs prepared with PC:Cer_16_ (90:10) were processed as in (a). **(e)** GUVs prepared with PC:Cer_brain_ (90:10) were processed as in (a). A total of 200-1050 GUVs were analyzed per condition in at least three independent experiments. PC (100), n=5; PC:Cer_16_ (90:10), n=3; PC:Cer_brain_ (90:10), n=4; PC:CL (90:10), n=5; PC:CL (90:10) + Bax, n=4. Data are means ± SD. Scale bar, 10 µm.

### Unlike cardiolipin, ceramide does not mediate Bax recruitment to model membranes

Previous work indicated that ceramide, generated in the mitochondrial outer membrane of mammalian cells upon irradiation, forms a platform into which Bax inserts, oligomerizes and functionalizes as a pore [30]. To determine whether ceramide directly facilitates insertion of Bax into membranes, we prepared membrane coated silica beads with liposomes composed of different lipid mixtures and monitored their ability to recruit DY-647P1-labeled Bax (Bax^647^) by confocal microscopy. Silica bead-supported lipid bilayers were visualized using lipophilic dyes DiI or DiO. Uncoated silica beads or silica beads coated with membranes composed of egg PC failed to recruit any appreciable amount of cBid-activated Bax^647^ (**Fig. 5a, c**). In contrast, cBid-activated Bax^647^ was readily and selectively mobilized to beads coated with membranes containing 10 mol% CL (**Fig. 5b, c**; **Fig. 6a**). As expected, omission of cBid treatment drastically reduced but did not eliminate CL-dependent membrane binding of Bax^647^ (**Fig. 5c; Suppl. Fig. 2**). These results are consistent with a crucial role of CL in promoting membrane recruitment of activated Bax and demonstrate that silica bead-supported lipid bilayers provide a suitable model for investigating the contribution of individual lipid species to membrane binding of Bax [41, 42]. When incubated with a mix of beads coated with lipid bilayers composed of either 10 mol% CL or Cer_16_ and 90 mol% egg PC, cBid-activated Bax^647^ was selectively recruited to the CL-containing membranes (**Fig. 6b, d**). In contrast, beads coated with Cer_16_-containing membranes failed to selectively mobilize cBid-activated Bax^647^ when mixed with beads coated with membranes composed of egg PC (**Fig. 6c, d**). These results indicate that Cer_16_, unlike CL, is unable to directly support membrane binding of cBid-activated Bax^647^.

**Figure 5.**
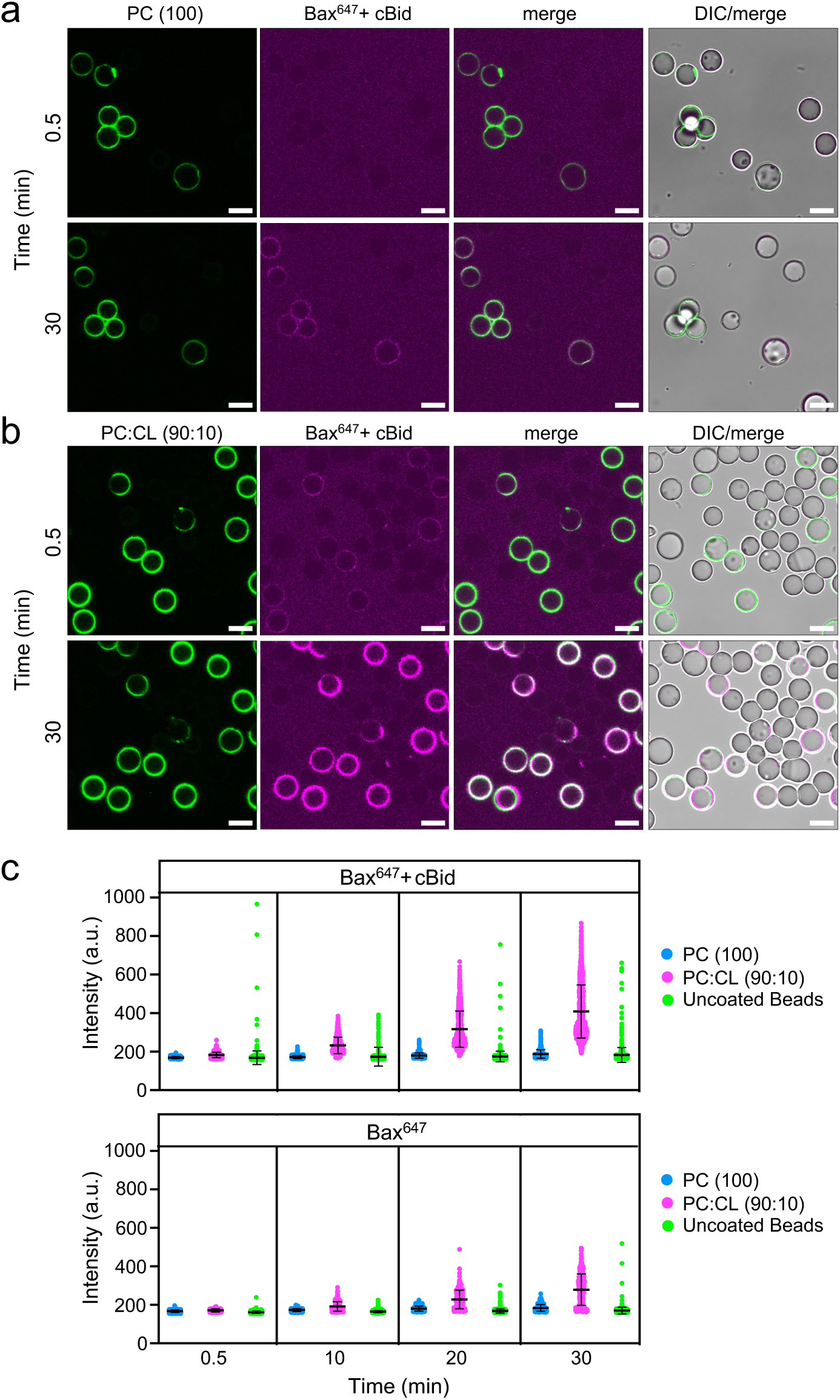
cBid-activated Bax is readily mobilized to cardiolipin-containing model membranes. **(a)** Silica beads coated with membranes prepared from PC (100; *green*) were mixed with uncoated beads and then incubated with 200 nM DY-647P1-labeled Bax (Bax^647^; *magenta*) and 50 nM cBid. At the indicated incubation times, beads were imaged by confocal fluorescence and differential interference contrast (DIC) microscopy. **(b)** Silica beads coated with membranes prepared from PC:CL (90:10; *green*) were mixed with uncoated beads and then incubated with 200 nM Bax^647^ (magenta) and 50 nM cBid. At the indicated incubation times, beads were imaged as in (a). Scale bar, 10 µm. **(c)** Bax^647^ bound to uncoated beads or beads coated with membranes prepared from PC (100) or PC:CL (90:10) was quantified at the indicated incubation times and expressed as relative fluorescence intensity. Data shown are based on a total of 600 to 900 individual measurements per condition in 3 independent experiments.

**Figure 6.**
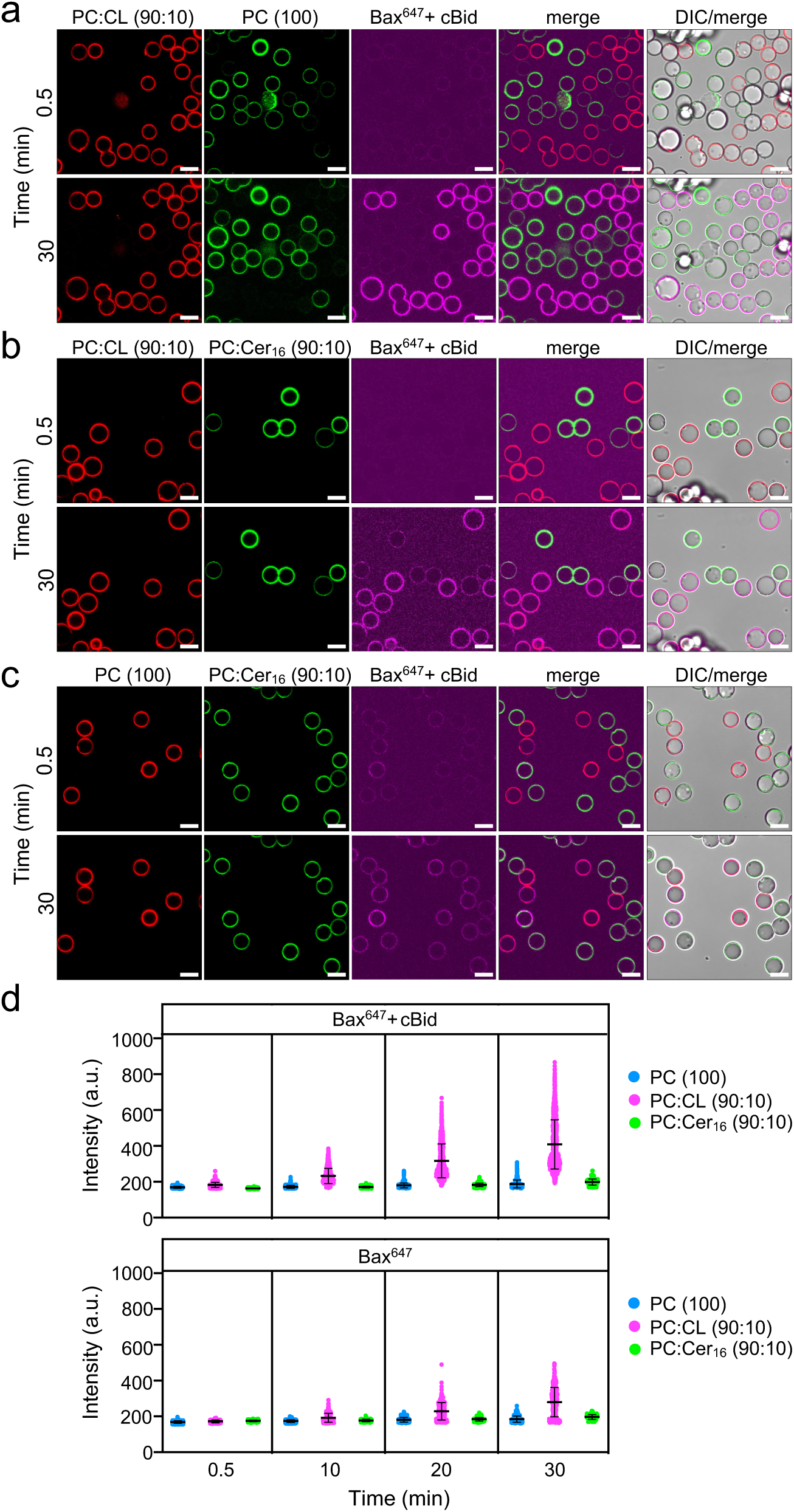
Ceramide cannot substitute for cardiolipin in mediating Bax recruitment to model membranes. **(a)** Silica beads coated with membranes prepared from PC (100; *red*) or PC:CL (90:10; *green*) were mixed and then incubated with 200 nM Bax^647^ (*magenta*) and 50 nM cBid. At the indicated incubation times, beads were imaged by confocal fluorescence and DIC microscopy. **(b)** Silica beads coated with membranes prepared from PC:CL (90:10; *red*) or PC:Cer_16_ (90:10; *green*) were mixed, incubated with Bax^647^ (*magenta*) and cBid, and then visualized as in (a). **(c)** Silica beads coated with membranes prepared from PC (100; *red*) or PC:Cer_16_ (90:10; *green*) were mixed, incubated with Bax^647^ (*magenta*) and cBid, and then visualized as in (a). Scale bar, 10 µm. **(d)** Bax^647^ bound to beads coated with membranes prepared from PC (100), PC:CL (90:10) or PC:Cer_16_ (90:10) was quantified at the indicated incubation times and expressed as relative fluorescence intensity. Data shown are based on a total of 350 to 900 individual measurements per condition in at least 3 independent experiments.

### Ceramide does not facilitate Bax-mediated pore assembly in model membranes

We next challenged a potential role of ceramide in facilitating the assembly of activated Bax in membrane permeabilizing pores. While addition of cBid-activated Bax to CL-containing GUVs led to their permeabilization for Dex10 and Dex70, substitution of CL for either egg PC or Cer_16_ completely abrogated the ability of Bax to form Dex10/Dex70-conducting pores (**Fig. 7**). Importantly, omission of cBid greatly reduced Bax-mediated pore formation in CL-containing GUVs (**Suppl. Fig. 3**) while cBid on its own did not influence GUV permeability for fluorescent dextrans (**Suppl. Fig. 4**). These results indicate that pore formation is strictly dependent on activated Bax and that ceramide cannot substitute for CL in facilitating formation of Bax pores that enable membrane passage of Dex10 and Dex70. To further validate this finding and explore a possible role for ceramide in the assembly of Bax pores with a smaller diameter, we next conducted calcein dequenching assays with CL- or ceramide-containing LUVs. Addition of cBid-activated Bax to LUVs containing 10 mol% CL readily triggered calcein release, which was abolished upon omission of cBid or Bax. When CL was substituted for egg PC, Cer_16_ or brain ceramide, cBid-activated Bax did not trigger any measurable calcein release (**Fig 8**). From this we conclude that ceramide is unable to replace CL in facilitating membrane binding, oligomerization and functionalization of Bax as a pore.

**Figure 7.**
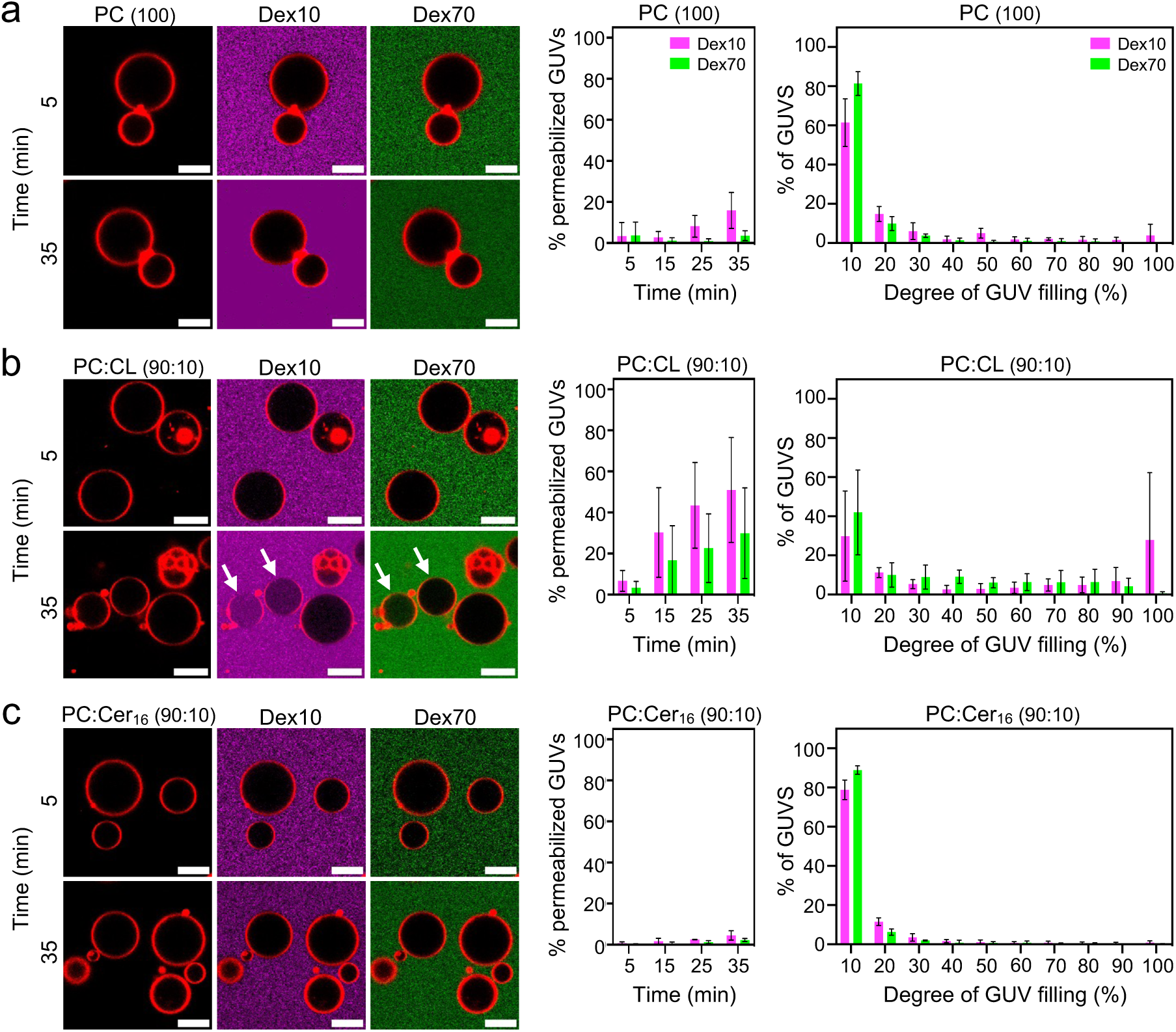
Ceramide cannot substitute for cardiolipin in supporting Bax-mediated pore formation. **(a)** GUVs prepared from PC (100) were stained with DiI (*red*) and incubated with 10 kDa dextran (Dex10; *magenta*) and 70 kDa dextran (Dex70; *green*) in the presence of 400 nM Bax and 50 nM cBid and then imaged using a LSM Airy scan microscope. At the indicated incubation time, the permeability of the GUVs for Dex10 and Dex70 was quantified. GUVs with a filling degree of 40% or more were considered permeable. In addition, the filling degree of the GUVs was determined at 35 min of incubation. **(b)** GUVs prepared with PC:CL (90:10) were processed as in (a). **(c)** GUVs prepared with PC:Cer_16_ (90:10) were processed as in (a). A total of 310-850 GUVs were analyzed per condition over at least three independent experiments. PC (100), n=4; PC:CL (90:10), n=4; PC:Cer_16_ (90:10). Data are means ± SD. Scale bar, 10 µm.

**Figure 8.**
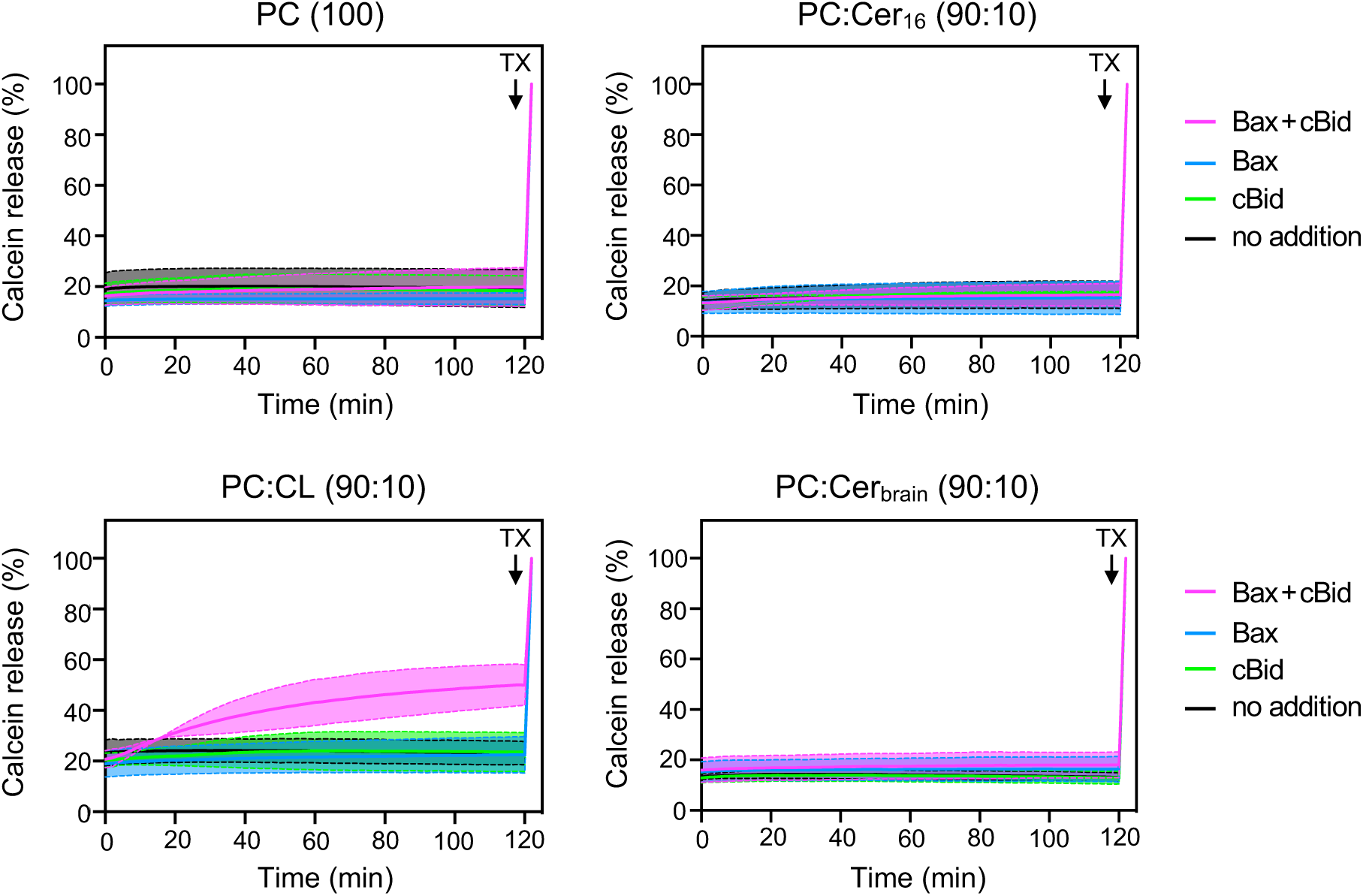
Ceramide does not facilitate Bax-mediated calcein release from vesicles. Real-time chromatograms of calcein released from PC:CL (80:20) vesicles incubated in the absence or presence of 100 nM Bax or 50 nM cBid. Leakiness of vesicles was tested by omitting Bax and cBid (no addition). Calcein release was measured over 120 min at 2 min intervals using a fluorescence plate reader. For each time point, the mean values of three independent measurements were determined and normalized to the value measured after Triton X100 (TX) addition, which served as a reference for 100% permeabilization.

## DISCUSSION

Compelling evidence indicates that ceramides can act directly on mitochondria to trigger apoptotic cell death but the underlying mechanism remains poorly understood [15, 45]. While one model postulates that ceramides form stable channels in the OMM to facilitate the release of cytochrome *c*, an alternative view is that ceramides self-assemble into microdomains that functionalize Bax to form cytochrome *c*-conducting pores [18, 19, 30]. Here, we challenged fundamental aspects of both models by analyzing the influence of ceramides on the leakiness of model membranes for a range of fluorescence size markers in the absence or presence of recombinant Bax. Our results indicate that ceramides on their own lack the propensity to permeabilize lipid bilayers for even the smallest fluorescence size markers. Moreover, we find that ceramides, unlike cardiolipin, neither facilitate membrane recruitment of Bax nor support the assembly of Bax into stable proteinaceous pores that enable passage of fluorescence markers in the size range of cytochrome *c*. Collectively, our data speak against a direct role of ceramides in apoptotic pore formation and indicate that the mechanism by which ceramides initiate permeabilization of the OMM is independent of ceramide channels or ceramide acting autonomously as translocation platform for Bax.

Ceramides have unique structural features that enable them to profoundly influence biophysical properties of membranes, promoting the formation of nonlamellar phases, inducing membrane negative curvature, and driving lateral domain segregation [46]. These bilayer-destabilizing properties of ceramides may be relevant for their ability to induce permeabilization of the OMM. Previous MD simulations [21], electrophysiological experiments on planar bilayers [18–20] as well as negative stain electron microscopy of ceramide-containing liposomes [24] indicated that ceramide-induced membrane permeabilization may occur through formation of stable ceramide-based channels with a diameter large enough to enable passage of cytochrome *c.* This apoptotic ceramide channel model was challenged in a study by Artetxe et al. [27]. These authors used a time-resolved liposome content leakage assay to monitor the release of calcein from sphingomyelin (SM)-containing vesicles upon SM-to-ceramide conversion by externally added SMase. They observed an overall gradual phenomenon of contents release instead of the all-or-none leakage expected for membrane permeabilization mediated by stable ceramide channels. In this experimental set-up, the observed contents leakage is likely due to membrane defects caused by a mismatch in the surface area between the two leaflets that originates from an asymmetric SM-to-ceramide conversion, as previously proposed [47, 48]. A disadvantage of the method used by Artetxe et al. is that the gradual contents leakage may have obscured all-or-none leakage events. To circumvent this limitation, we here analyzed the leakiness of GUVs containing 10 mol% Cer_16_ or brain ceramides for externally added fluorescence size markers. This revealed that inclusion of either source of ceramides did not affect the permeability of GUVs for fluorescence markers in the size range of cytochrome *c*. Our results complement the findings by Artetxe et al. and argue against ceramide channels as the primary mechanism by which ceramides initiate permeabilization of the OMM to commit cells to death.

Our data recapitulate a crucial role of cardiolipin (CL) in supporting membrane binding, insertion and permeabilization of cBid-activated Bax, as documented previously in numerous other studies [39, 49–51]. CL is a unique dimeric phospholipid that exists almost exclusively in the inner mitochondrial membrane (IMM). The main structural feature that distinguishes CL from other common phospholipids is the fact that a single headgroup alcohol, a glycerol molecule, is shared by two phosphatidate moieties. This arrangement has important implications regarding the physical properties of CL within the context of a lipid bilayer [52]. The reduced effective cross-sectional area of the CL polar headgroup relative to that of the four hydrocarbon chains enhances its propensity to form inverted nonlamellar phases [53]. Due to the small size and impaired flexibility of the CL headgroup, its capacity for steric self-shielding of the charged phosphate groups is diminished. This makes CL as component of cell membranes more “exposed” and prone to specific interactions with proteins. These properties likely enable CL to serve as chaperone in facilitating Bax-insertion, oligomerization and pore opening [43, 54, 55]. Even though ceramides and CL have overlapping bilayer-destabilizing properties, our findings clearly indicate that ceramides on their own neither support membrane binding of cBid-activated Bax, nor facilitate Bax-mediated pore assembly in model membranes. These results are hard to reconcile with the idea that ceramides generated in the OMM upon radiation of mammalian cells autonomously form microdomains into which Bax inserts and oligomerizes into a cytochrome *c*-conducting pore [56].

We previously reported that diverting CERT-mediated ceramide transport to mitochondria triggers Bax-dependent apoptosis, that this process requires ongoing *de novo* ceramide synthesis, and that apoptosis induction relies on intact ceramides rather than downstream intermediates of ceramide turnover [15, 16]. A subsequent chemical screen for ceramide binding proteins combined with computer simulations and functional studies in cancer cells led to identification of the OMM channel proteins VDAC1 and VDAC2 as core components of a potential pathway through which ceramides may exert their pro-apoptotic activities [57]. Other work revealed that VDAC2 specifies Bax recruitment to mitochondria and controls the pro-apoptotic activity of Bax by mediating its retro-translocation into the cytosol [58]. Interestingly, VDAC residues involved in ceramide binding also directly participate in mobilizing hexokinase-I (HKI) to mitochondria [59], a condition thought to promote cell growth and survival in hyperglycolytic tumors [60–62]. Together, these results point at an alternative mechanism by which mitochondrial ceramides may initiate OMM permeabilization and apoptotic cell death, namely as modulators of VDAC-based translocation platforms for pro-apoptotic Bax and anti-apoptotic HKI. A challenging but desirable prospect is to experimentally challenge this model.

## METHODS

### Chemicals and antibodies

L-α-phosphatidylcholine (egg PC; cat. no. 840051P), heart cardiolipin (CL; cat. no. 840012C), brain ceramide (Cer_brain_; N-octadecanoyl-D-erythro-sphingosineN-(octadecanoyl)-sphing-4-enine; cat. no. 8600052P) and C16:0 ceramide (Cer_16_; N-palmitoyl-D-erythro-sphingosine; cat. no. 860515) were purchased from Avanti Polar Lipids. The lipophilic dyes DiI (cat. no. D3911) and DiO (cat. no. D275) were from Invitrogen. The protein dyes DY-547P1 (cat. no. 547P1-03) and DY-647P1 (cat. no. 647P1-03) were from Dyomics. Calcein (cat. no. C0875) was from Merck. Dextran Alexa Fluor 647 10 kDa (cat. no. D22914) and Dextran fluorescein 70 kDa (cat. no. D1823) were from Invitrogen. cBid (cat. no. 882-B8) was from R&D Systems. α-Bax rabbit monoclonal antibody (cat. no. 5023s) was from Cell signaling, α-rabbit HRP (cat. no. 170-6515) was from BioRad. Chemicals for buffers were purchased from AppliChem, Carl Roth or Sigma Aldrich. All aqueous solutions were prepared using ultrapure water, degassed and filtered through 0.2 μm hydrophilic membranes

### Purification and cysteine labelling of recombinant Bax

The bacterial expression construct encoding the human Bax single cysteine mutant S4C/C62S/C126S tagged with a *C*-terminal intein–chitin binding domain (CBD) was described in [63] and used to transform the *E. coli* expression strain BL21 codonplus (DE3)-RIL. Transformants were grown in terrific broth [1] medium (24 g/l yeast extract, 12 g/l Casein, 9,4 g/l K_2_HPO_4_, 2,2 g/l KH_2_PO_4_, pH 7.2) containing ampicillin (100 µg/ml) and chloramphenicol (34 µg/ml) at 37°C while shaking. Bax expression was induced at an OD_600_ of 0.4-0.6 by adding IPTG to a final concentration of 1 mM followed by incubation overnight at 20°C while shaking. Cells were harvested by centrifugation (5,350 x g, 15 min, 4°C), resuspended in Chitin Buffer (CB; 500 mM NaCl, 20 mM Tris pH 8.1) containing ∼0.2 mg/mL DNAseI and 1 x Protease Inhibitor Cocktail (PIC, 1 mg/ml apoprotein,1 mg/ml leupeptin,1 mg/ml pepstatin, 5 mg/ml antipain, 157 mg/ml benzamidine), and lysed using a high-pressure cell disrupter (Basic Z) with a 0.18 mm tip and disruption pressure of 0.45 kBar. After centrifugation to remove debris (14,000 x g, 1 h, 4°C), the lysate was incubated with chitin-affinity resin (New England Biolabs; cat. no. S6651) pre-equilibrated with CB for 2 h at 4°C while rotating. After washing the resin with 10 column volumes (CV) of CB, intein self-cleavage was induced by equilibrating the matrix with 3 CV of CB containing 50 mM DTT followed by an overnight incubation in 1 CV of the same buffer. Bax was collected in the flow through after applying 2 CV of CB, dialyzed against Mono A buffer (20 mM Tris pH 8.0) for 4h with buffer exchanges every h, and loaded onto an anion exchange column (1 ml Fractogel EMB TMAE High Cap M; Merck; cat. no. 1.10316.0100). After washing the column with 30 CV of Mono A buffer, a linear salt gradient (0-1M NaCl) was applied in Mono A buffer at a flow rate 0.5 mL/min. Bax peak fractions were pooled and 35 µM of the protein was subjected to cysteine labeling with 105 µM DY-647P1 (Dyomics) for 45 min at RT in the dark. After addition of 315 µM of L-Cysteine, the reaction mixture was incubated for 15 min at RT, loaded onto a size exclusion chromatography column (Superdex 200 Increase 10/300 GL column; Cytiva; cat. no. 28990944) and in SEC Buffer (20 mM Tris, 150 mM NaCl pH 8.0). Protein purity and efficiency of cysteine labeling was determined by SDS-PAGE, colloid Coomassie Blue staining and in-gel fluorescence (IGF) analysis using a Typhoon FLA 9500 Biomolecular Imager (GE Healthcare Life Sciences). Immunoblot analysis was performed using α-Bax rabbit monoclonal antibody (1:1000) and α-rabbit HRP antibody (1:5000) and immunoreactive bands were imaged using a Gel Doc XRS scanner (BioRad). Gels and immunoblot images were processed with ImageLab software (BioRad). Purified Bax was snap-froozen and stored at −80°C at a concentration of 10 µM for the unlabeled protein and at 6.25 µM for the DY-647P1-labeled protein (labeling efficiency: ∼74%).

### Calcein release assay

Lipid mixtures were prepared in chloroform and dried in a vacuum desiccator to create a thin lipid film. Lipid mixtures used were PC (100 mol%), PC:CL (80:20 or 90:10 mol%), PC:Cer_16_ (90:10 mol%) and PC:Cer_brain_ (90:10 mol%). The lipid film was resuspended in 80 mM calcein (dissolved in ultrapure H_2_O, pH 7.5) by vigorous vortexing to a final concentration of 4 mg/mL and subjected to five freeze-thaw cycles using liquid nitrogen and a 37°C water bath. To obtain large unilamellar vesicles (LUVs), the lipid suspension was extruded 11 times through a 0.4 μm track-etched polycarbonate membrane (Whatman-Nuclepore) and then 21 times through a 0.1 μm polycarbon membrane using a mini-extruder (Avanti Polar Lipids). Free calcein was removed by applying the liposome mixture onto a G25 fine Sephadex matrix (Cytiva; cat. no. 17003201) pre-equilibrated with Outside Buffer (OB; 140 mM NaCl, 20 mM Hepes, 1 mM EDTA, pH 7.0). The quality of calcein-loaded LUVs was verified by measuring calcein fluorescence before and after addition of 0.25% Triton X100 using a fluorescence plate reader (Tecan Infinite M). Samples were excited at 490 nm and emission was measured at 520 nm. For measuring Bax/cBid-dependent membrane permeabilization, Bax (100 nM) and cBid (50 nM) were added in 100 μL OB per well of a 96-well plate that was pre-blocked with BSA. After addition of calcein-loaded LUVs to a final dilution of 1:40, calcein fluorescence intensity was measured every 2 min for 2 h at RT. At the end of the reaction, 0.25 % Triton X100 was added to each well and fluorescence was measured again to determine maximal calcein release. For each time point, the mean values of three independent measurements were determined and normalized to the value measured after Triton X100 addition, which served as a reference for 100% permeabilization.

### Membrane-coated silica beads

Membrane-coated silica beads were produced as described in Maib and Murray [42] with some modifications. Lipid films were resuspended in Buffer 1 (200 mM NaCl) to a final concentration of 1 mg/mL, subjected to five freeze thaw cycles, and then extruded through 0.4 μm and 0.1 μm track-etched polycarbonate membranes as described above. Silica beads (10 μm; Whitehouse Scientific) were washed in EtOH, H_2_O and Buffer 1. Membrane-coated silica beads were generated by mixing 0.2 mg of silica beads with 10 µg liposomes containing 0.01 mol% DiI or DiO in 100 µl Buffer 1. To enable proper membrane coating, the mixture was incubated for 45 min on ice. Beads were washed twice in 500 µl Buffer 2 (200 mM Hepes, pH 7.0, blocked with 1 mg/ml BSA in Buffer 3 (150 mM NaCl, 20 mM Hepes pH 7.0) for 30 min at RT, and then resuspended in 300 µl Buffer 3. To assess Bax binding, 50 µl of the bead suspension was added to uncoated m-Slide 8 well chambers (Ibidi). DY-647P1-labeled Bax (Bax^647^) was pre-mixed with cBid in Buffer 3 and then added dropwise to the beads at a final concentration of 200 nM and 50 nM, respectively. This was followed by immediate imaging of the beads at RT for a period of 30 min with 10 min intervals. Confocal images were acquired with an inverted Olympus IX83-P2ZF LSM microscope equipped with a sCMOS camera (Hamamatsu ORCAFlash 4.0 V3) and a Coherent OBIS LX/LS laser system with the following wavelengths: 488 (20 mW), 561 (20 mW) and 640 (40 mW). Images were captured using an oil immersion PLAPON-SC 60x super-corrected apochromatic objective with a NA of 1.40. For precise positioning of the beads, a motorized xy-stage (IX3-SSU, Olympus) was used. Confocal point scanning was enabled using a xy-piezo stage (P-542.2SL, Physik Instrumente). Bax^647^ binding was quantified using an ImageJ script adapted from Maib and Murray [42] and provided in Supplementary Information. The signal from DiO or DiI was used to segment each bead individually, thereby creating a mask that was used to measure Bax^647^ recruitment to each bead. Beads with a perturbed membrane coating were excluded from the analysis. Bax^647^ signals were plotted using GraphPad Prism software version 8.3.0 (GraphPad Software LLC, USA).

### GUV permeabilization assay

For GUV formation, the following lipid mixtures were prepared in chloroform: PC (100 mol%), PC:CL (80:20 or 90:10 mol%), PC:Cer_16_ (90:10 mol%) and PC:Cer_brain_ (90:10 mol%). After addition of 0.01 mol% DiI, the lipid mixtures were added dropwise on the conductive site of an indium tin oxide (ITO) glass. In total 40 µg lipid was applied and then dried for 30 min at RT in a vacuum desiccator (Nalgene). The lipid containing ITO glass was sandwiched with a clean ITO glass using 3 mm-thick Teflon spacers to create a chamber that was filled with 220 µl sucrose (300 mM). GUVs were formed by applying an AC electric field (10 Hz, 1.5 V amplitude) for 60 min at RT and deattached from the glass surface by changing the frequency to 2 Hz for 30 min. The GUVs were then gently transferred into an eppendorf tube containing 400 µl PBS and incubated for 20 min at RT. To measure GUV permeabilization, fluorescein-labeled 70 kDa dextran and Alexa Fluor647-labeled 10 kDa dextran were added to m-Slide 8 well chambers (Ibidi) pre-blocked with BSA in PBS at a final concentration of 2.5 µM each. Bax and cBid were added to a final concentration of 200 nM and 50 nM, respectively. In the final step, 50 µl of the GUV suspension collected from the bottom of the eppendorf tube were added to reach a final reaction volume of 200 µl per well. After 5 min of incubation, GUV permeabilization was monitored for 30 min with 10 min intervals using an LSM 880 Airy scan microscope (Zeiss cell observer 7) and a Pln-Apochromat 63 X oil immersion objective (Zeiss) with a NA of 1.4. Excitation light came from argon ion (458 nm, 488 nm, 514 nm), Diode 405-30 (405 nm), DPSS 561-10 (561 nm), or HeNe lasers (633 nm). A spectral beam guide was used to separate emitted fluorescence. GUVs were positioned using a xy scanning stage SCAN IM 130×100 (Märzhäuser) and a high precision z-axis piezo stage (Wienecke & Sienske). GUV permeabilization was quantified using GUV detector software from the University of Tübingen [64]. The degree of GUV filling was calculated using the following formula:

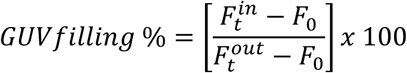

with Ft^in^ and Ft^out^ signifying the average fluorescence intensities in- and outside of a GUV at a time t. F_0_ is the background fluorescence.

## ACKNOWLEDGEMENTS

This work was supported by the Deutsche Forschungsgemeinschaft (378148610, 448344643 and 467522186/SFB1557–P9 to J. C. M. H.; 467522186/SFB1557–P5 to K.C.).

## Author Contributions

J. C. M. H. designed the research; M. W. performed experiments with critical input from S. K., and B. F.; K. C. provided expertise on membrane permeabilization assays and experiments with recombinant Bax, and helped interpret the data; M. W. and J. C. M. H. wrote the manuscript; all authors discussed results and commented on the manuscript.

## Data Availability

All data generated or analyzed in this study are included in the manuscript and supporting files. Source data with sample sizes, number of technical and/or biological replicates, means, standard deviations, and calculated *p* values (where applicable) are provided in the Supplementary Data file. Uncropped scans of immunoblots are provided in Supplementary Information.

## Data availability

All data generated in this study are included in the manuscript and Supplementary Information file.

