## Supplemental Information for "Challenging a role for ceramide channels and microdomains in apoptosis induction using a bottom-up approach"

**This PDF file includes:**

Supplementary Figs. 1 to 4

ImageJ Macro „Silica Beads Intensity“

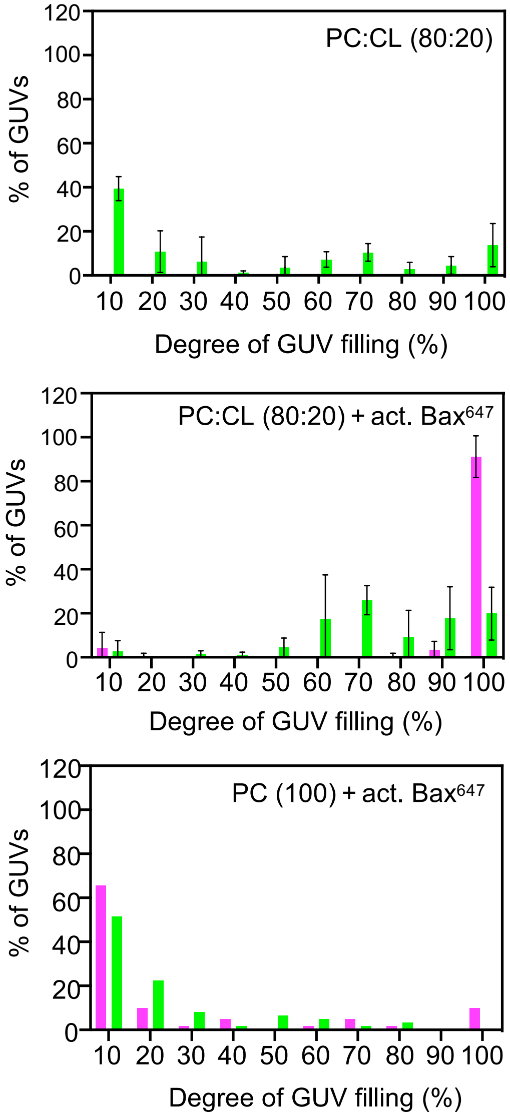


**Suppl. Figure 1. Quantification of Bax-induced permeabilization of cardiolipin-containing GUVs**

GUVs prepared from PC (100) or PC:CL (80:20) were incubated with calcein in the absence or presence of 400 nM $Bax^{647}$. After 35 min, the distribution of the filling degree of GUVs with calcein and $Bax^{647}$ was determined. GUVs with a degree of filling of 40 % or more were considered permeable. For GUVs prepared from PC:CL (80:20), a total number of 70-100 GUVs were analyzed in three independent experiments. For GUVs prepared from PC (100), a total of 50 GUVs were analyzed in one experiment. Data are means ± SD.

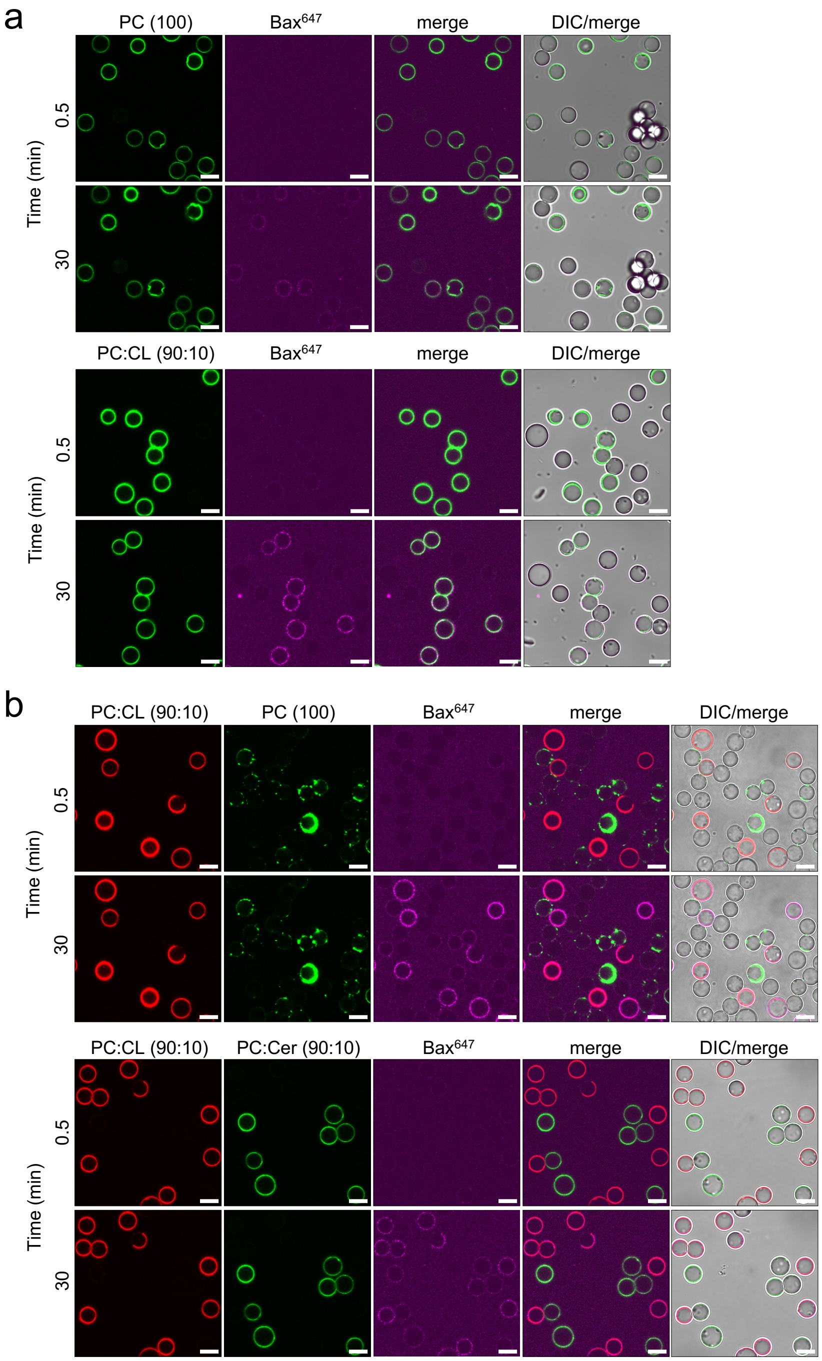


**Suppl. Figure 2. Recruitment of Bax to cardiolipin-containing model membranes is not strictly dependent on cBid activation**

(**a**) Silica beads coated with membranes prepared from PC (100; *green*) or PC:CL (80:20; *green*) were mixed with uncoated beads and incubated with 200 nM DY-647P1-labeled Bax ($Bax^{647}$; *magenta*). At the indicated incubation times, beads were imaged by confocal fluorescence and differential interference contrast (DIC) microscopy.

(**b**) Silica beads coated with membranes prepared from PC:CL (90:10; *red*) and PC (100; *green*) or $PC:Cer_{16}$ (90:10; *green*) were mixed and then incubated with 200 nM $Bax^{647}$ (*magenta*). At the indicated incubation times, beads were imaged as in (a). Scale bar, 10 µm.

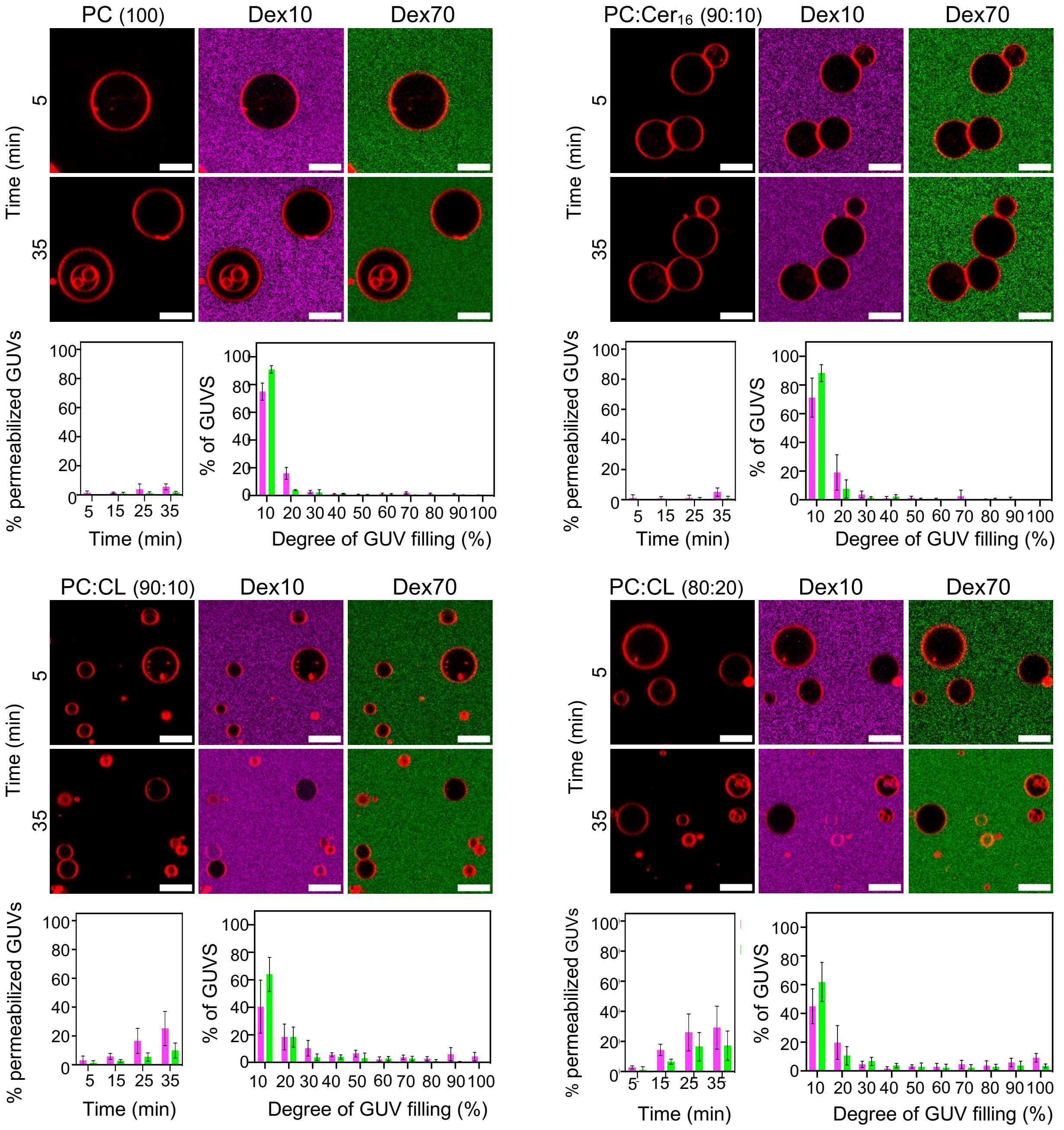


**Suppl. Figure 3. Bax-mediated pore formation critically relies on cardiolipin and cBid**
GUVs prepared from PC (100), PC:Cer$_{16}$ (90:10), PC:CL (90:10), or PC:CL (80:20) were stained with DiI (*red*) and incubated with 10 kDa dextran (Dex10; *magenta*) and 70 kDa dextran (Dex70; *green*) in the presence of 200 nM Bax. GUVs were imaged by LSM Airy scan microscopy. At the indicated incubation times, the permeability of GUVs for Dex10 (*magenta*) and Dex70 (*green*) was determined. GUVs with a filling degree of 40 % or more were considered permeable. The distribution of the filling degree of GUVs with Dex10 and Dex70 was determined at 35 min of incubation. In total 225-570 GUVs were analyzed per condition from at least three experiments. PC (100), n=3; PC:Cer$_{16}$ (90:10), n=3; PC:CL (90:10), n=4; PC:CL (80:30), n=3. Data are means ± SD. Scale bar, 10 µm.

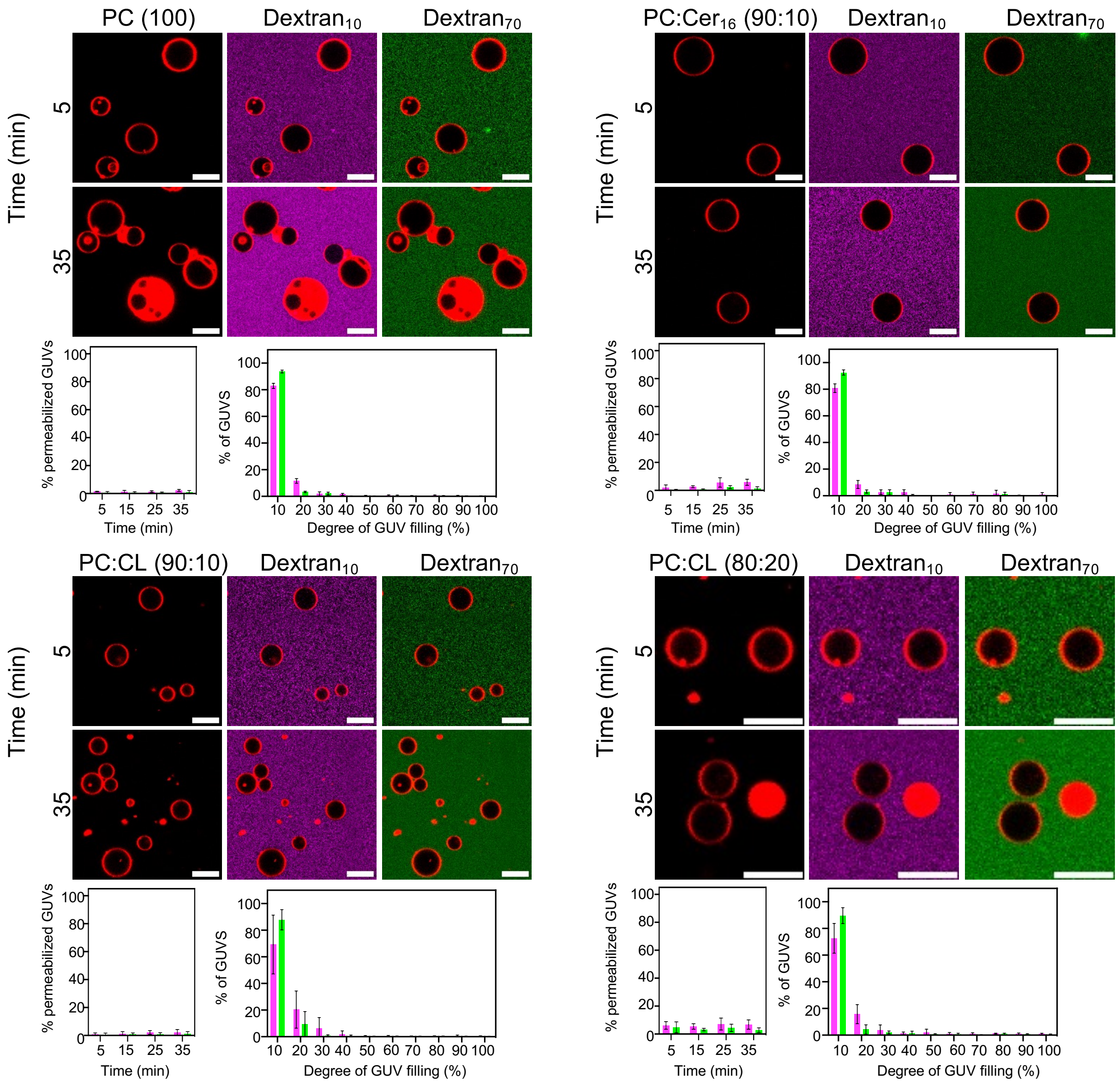


**Suppl. Figure 4. cBid lacks pore forming activity**

GUVs prepared from PC (100), PC:$Cer_{16}$ (90:10), PC:CL (90:10), or PC:CL (80:20) were stained with DiI (*red*) and incubated with 10 kDa dextran (Dex10; *magenta*) and 70kDa dextran (Dex70; *green*) in the presence of 50 nM cBid. GUVs were imaged by LSM Airy scan microscopy. At the indicated incubation times, the permeability of GUVs for Dex10 (*magenta*) and Dex70 (*green*) was determined. GUVs with a filling degree of 40 % or more were considered permeable. The distribution of the filling degree of GUVs with Dex10 and Dex70 was determined at 35 min of incubation. In total 220-1050 GUVs were analyzed per condition from at least three experiments. PC (100), n=3; PC:$Cer_{16}$ (90:10), n=3; PC:CL (90:10), n=4; PC:CL (80:30), n=3. Data are means ± SD. Scale bar, 10 µm.

ImageJ Macro „Silica Beads Intensity“

*>>>Start of code*

```
if (File.exists(getDirectory("plugins")+"/Read_and_Write_Excel-1.1.7.jar") != 1){ //checking for
the excel plugin. If there are updates, also update the version number here
	exit("ResultsToExcel Plugin not found. Make sure you activated the plugin in the ImageJ
updater. ('Help' > 'Update...' > 'Manage update sites')'");
}

dir1 = getDirectory("Choose Source Directory ");
list = getFileList(dir1);
run("Read and Write Excel", "file_mode=read_and_open
file=["+dir1+"/BeadsMacroResults.xlsx] sheet=[Bax to DiO]");

for (i=0; i < list.length; i++) {
	filename = dir1 + list[i];

	if (File.isDirectory(filename) != 1 && endsWith(filename, "oir") == 1) { //check if
current object is not a folder and is a correct file type
		run("Bio-Formats Importer", "open=["+filename+"] autoscale
color_mode=Default rois_import=[ROI manager] view=Hyperstack stack_order=XYCZT");

		name = getTitle(); //this get the current title of the image as variable
		selectWindow(name); //this just tells ImageJ to select the current image
		run("Split Channels"); //this just split the channels and seperate them

		//HW: Data analysis with first channel as reference:
		selectWindow("C1-"+name); //this selects the first lipid mask for segmentation. in
this case the DiO signal
		run("Duplicate...", "title=C1-double.oir");
		selectWindow("C1-"+name);
		setAutoThreshold("Li dark"); //this segemnts the first channel to use as mask.
Here you will have to play around to find the best segementation option for your needs
		run("Convert to Mask", "method=Li background=Dark calculate");
		setOption("BlackBackground", false);
		run("Fill Holes"); //this just closes small holes resulting from segemntation errors
		run("Watershed"); //this seperates beads that are close to each other into
individual beads
		run("Analyze Particles...", "size=15-250 circularity=0.2-1.00 show=Overlay clear
add stack"); //this is important. here the size and circularity of the segemntation is used to detect
the beads. Depending on your imaging settings, the size needs to be adjusted!
		//HW: the parameter "clear" leads to clearing the ROI manager, which was
lacking and lead to reading out ROIs of former analysis
```

```
		upperROI = roiManager("count"); //this whole loop is really the smart part about
the macro. It creates a small ring around your beads and only detects the signal that is bound to
the beads. If you want to change the thickness of the ring, adjust the enlarge parameters. It does
so for everyparticle that has been detected previously.
	for (index = 0; index < upperROI; index++) {
	roiManager("Select", index);
	run("Enlarge...", "enlarge=-1");
	run("Make Band...", "band=1.25");
	roiManager("Update");
	}

		roiManager("Deselect"); //from here the user part begins
		selectWindow("C1-"+name);
		setLocation(screenWidth/2-getWidth()*getZoom(), 100);
		selectWindow("C1-double.oir");
		setLocation(screenWidth/2, 100);
		selectWindow("ROI Manager");
		setLocation(screenWidth/2, screenHeight/2);
		waitForUser("Evaluate ROIs", "Delete bad ROIs in the ROI-Manager by selecting
clicking 'Delete'. After that, click OK to proceed.");

		selectWindow("C3-"+name); //here it now selects the window for the
measruments. It is important that the correct channel is selected. In this case channel three with
the Bax647 probe
		roiManager("Deselect");
		roiManager("Measure");
		run("Read and Write Excel", "file_mode=queue_write no_count_column
dataset_label=["+name+"] sheet=[Bax to DiO]"); //remove "stack_results" to write them next to
each other
		run("Clear Results");

		//HW: Data analysis with second channel as reference:
		selectWindow("C2-"+name); //this selects the first lipid mask for segmentation. in
this case the DiI signal
		run("Duplicate...", "title=C2-double.oir");
		selectWindow("C2-"+name);
		setAutoThreshold("Li dark"); //this segemnts the first channel to use as mask.
Here you will have to play around to find the best segementation option for your needs
		run("Convert to Mask", "method=Li background=Dark calculate");
		setOption("BlackBackground", false);
		run("Fill Holes"); //this just closes small holes resulting from segemntation errors
		run("Watershed"); //this seperates beads that are close to each other into
individual beads
		run("Analyze Particles...", "size=10-650 circularity=0.05-1.00 show=Overlay
clear add stack"); //this is important. here the size and circularity of the segemntation is used to
detect the beads. Depending on your imaging settings, the size needs to be adjusted!
```

```
		upperROI = roiManager("count"); //this whole loop is really the smart part about
the macro. It creates a small ring around your beads and only detects the signal that is bound to
the beads. If you want to change the thickness of the ring, adjust the enlarge parameters. It does
so for everyparticle that has been detected previously.
	for (index = 0; index<upperROI; index++) {
	roiManager("Select", index);
		run("Enlarge...", "enlarge=-1");
	run("Make Band...", "band=1.25");
	roiManager("Update");
	}

		roiManager("Deselect"); //from here the user part begins
		selectWindow("C2-"+name);
		setLocation(screenWidth/2-getWidth()*getZoom(), 100);
		selectWindow("C2-double.oir");
		setLocation(screenWidth/2, 100);
		selectWindow("ROI Manager");
		setLocation(screenWidth/2, screenHeight/2);
		waitForUser("Evaluate ROIs", "Delete bad ROIs in the ROI-Manager by selecting
clicking 'Delete'. After that, click OK to proceed.");

		selectWindow("C3-"+name); //here it now selects the window for the
measruments. It is important that the correct channel is selected. In this case channel 3 with the
Bax647 probe
		roiManager("Deselect");
		roiManager("Measure");

		run("Read and Write Excel", "file_mode=queue_write no_count_column
dataset_label=["+name+"] sheet=[Bax to DiI]"); //remove "stack_results" to write them next to
each other
		run("Close All");
		close("Results");
		close("ROI Manager");
	}
}
run("Read and Write Excel", "file_mode=write_and_close");
waitForUser("Finish", "Script finished");
```

*>>>End of code*
